# Effect of calcium intake and dietary cation anion difference in early lactation on bone mobilization dynamics all over lactation in dairy cows

**DOI:** 10.1101/674184

**Authors:** Pierre. Gaignon, Karine Le Grand, Anca-Lucia Laza-Knoerr, Catherine Hurtaud, Anne Boudon

## Abstract

This study aimed to evaluate the consequences of increased bone mobilization in early lactation on the dynamics of the milk Ca content during lactation and bone reconstitution. Fifteen multiparous Holstein cows were distributed among 3 treatments 5 weeks before their expected calving date. Those treatments consisted of the provision of dedicated diets through the first 10 weeks of lactation. During that period, the control treatment (NCa) consisted of a diet providing 100\% of the Ca requirements, with a dietary cation-anion difference (DCAD) of 200 mEq/kg DM. The treatments LCa (Low Ca) and LCaLD (Low Ca, Low DCAD) consisted of diets providing 70% of the Ca requirements, with a DCAD of 200 and 0 mEq/kg DM for LCa and LCaLD, respectively. After 10 weeks, all cows received the same total mixed ration which was formulated to meet 100\% of the Ca requirements. LCa and LCaLD tended to decrease the body retention of Ca at 3 weeks of lactation compared with NCa, but did not affect either the dynamics of the blood biomarkers of bone formation and resorption during the lactation or the body retention of Ca at 17 weeks of lactation. Cows almost entirely compensated for the decrease in Ca supply by increasing their apparent digestive absorption of Ca at 3 weeks of lactation, whereas the apparent digestive absorption was unaffected by the treatments at 17 weeks of lactation. Milk production tended to be lower throughout lactation with the LCa and LCaLD compared with the NCa, with a mean difference of 2 kg/d. This study indicated that measuring the dynamics of the milk Ca content during lactation cannot be considered effective for indirectly estimating the dynamics of bone mobilization of cows. The results also showed that limited Ca intake at the beginning of lactation can have deleterious effect on milk production.

## Introduction

Dairy cows excrete a large amount of Ca during lactation due to the high content of Ca in milk (1), and this Ca flow suddenly and importantly increases later in lactation (2). During the first months of lactation, the dietary Ca intake is generally lower than the amount of Ca excreted in the milk, feces or urine (1), and several responses occur in the face of this imbalance. The first response is an increase in bone resorption, mediated by the secretion of 2 hormones, PTH and PTHrP, which allow the use, by other organs, of the Ca contained in the mineralized matrix of bone (3). The net mobilization of bone resulting from this increase in bone resorption at the beginning of lactation can reach up to 10 to 20% of the bone mass during lactation, with the amount of mobilized Ca being replenished later in lactation (4–6). The other responses, such as an increase in the intestinal Ca absorption or a decrease in urinary loss, occur later (7,8), explaining the existence of cycles of bone mobilization and reconstitution during lactation in cows (9–13).

Questions remain about the consequences of the amplitude and the completeness of those cycles of bone mobilization and reconstitution on the health and productivity of dairy cows. Incomplete bone reconstitution at the end of the lactation can result in a higher susceptibility of cows to the restricted supply of P during the following lactation, with suboptimal production performances, as highlighted for sucker cows by Dixon et al. (6), or maybe to higher susceptibility of milk fever at the beginning of the next lactation (14). The effects of dietary Ca and P content and strategies for the supplementation of Ca and P on the amplitude and the completeness of the cycle of bone formation and resorption have been quantified in several experiments in lactating ruminants (10–13,15,16).

Confirmation of an effect of the amplitude and completeness of the bone cycles on the health and productivity of dairy cows would consequently affect the estimations of the Ca and P requirements of those animals. Current recommendations are based on the principle that daily excretions of Ca and P allow a certain production level, with minimal fecal and urinary losses and with replacement by an equivalent amount of daily intake of those elements (17–19). This principle does not consider that bone mobilization in early lactation and the reconstitution in late lactation that could constitute either an extra-supply or a specific requirement. Possibly, a strategy of supplementation would have to be determined at the scale of the whole cycle of lactation-gestation, by taking into account bone reserves. However, the amount of published research available remains too limited to allow a definition of an optimal supplementation strategy of Ca and P at this scale (17–19).

A major limit to addressing these issues is the lack of fast and cost-efficient methods for evaluating, in a significant number of cows, the amplitude and eventually the completeness of the cycles of bone mobilization and reconstitution during lactation and gestation. Measurements of retained Ca and P at the animal scale (11) allow an evaluation of the daily net flow of mobilized and reconstituted Ca and P reserves, but it is very time-consuming and requires keeping the animals individually stabled. Bone biopsies (9) allow a good evaluation of bone reserve of Ca and P, but the number of repetitions of the measurement per animal cannot be multiplied on an significant number of animals because of the time, cost and ethics. The concomitant use of blood biomarker analyses of bone formation and resorption increased during the last 20 years, and this interesting method allows monitoring of the relative dynamics of bone formation and resorption during lactation (13,20,21). This method can be applied to a relatively high number of cows but is limited by the necessity for conducting several blood samplings during lactation, with relatively expensive analyses.

VanHouten et al. (22) showed that a decrease in Ca intake in mice induced a lower Ca secretion in milk and a higher bone resorption mediated by PTHrP secretion, with both mediated by the Ca-sensing receptor (**CaSR**) in the mammary gland; those results suggest that the monitoring of milk Ca content during lactation could be an inexpensive way to indirectly estimate the dynamics of bone resorption. Mid-infrared spectra technology allows a rapid and inexpensive way to determine the milk Ca content (23,24). Data collected during several stages of lactation in dairy cows with different parities (25) suggested that milk Ca and P contents could be related to the plasma concentrations of biomarker of bone formation and resorption. However, whether those variations are specifically related to the cycle of bone formation and mobilization or other interfering effects is unknown. Thus, the objective of this experiment was to induce bone mobilization in lactating cows through dietary treatments and to determine the consequences on i) the dynamics of milk Ca and P contents, of the plasma concentrations of biomarkers of bone formation and resorption, and the retention of Ca and P by the body, and ii) on bone reconstitution dynamics in late lactation. The dietary treatments either supplied Ca and P according to the French recommendation (19), restricted the Ca supply, or restricted the Ca supply and decreased the dietary anion cation difference (DCAD) known to increase bone resorption. Our hypothesis was that the restricted Ca supply, with or without low DCAD, would increase bone mobilization in early lactation and decrease the milk Ca content during this period. The increase in bone mobilization was expected to be clearer with the low DCAD diet. Another hypothesis was that the increased bone mobilization in early lactation would induce a higher bone reconstitution in late lactation.

## Material and Methods

### Animals and Experimental Design

The 3 compared treatments consisted of 3 mineral supplementations fed between 5 d after calving and 10 weeks from the beginning of lactation. The Ca content of the supplement was either calculated to allow a fully meet the Ca requirements of the cows according to the INRA feeding system (19), with a DCAD of 200 mEq/kg DM (Normal Ca, **NCa**), or it was calculated to provide only 70% of the Ca requirements of the cows, with a DCAD of either 200 (Low Ca, **LCa**) or 0 (Low Ca and Low DCAD, **LCaLD**) mEq/kg DM. These 3 treatments were planned to allow comparisons among the 15 lactating cows according to a complete randomized block design, with lactation stage as the blocking factor. Five weeks before the calving date of the cow that was expected to calve first, 18 multiparous Holstein cows were blocked into three groups of 6 cows according to their expected dates of calving. Cows were assigned to the three treatments to allow a homogenous representation of groups and parity within each treatment, and as similar as possible, similar averages of mature equivalent milk production and milk protein contents as observed in the first 32 weeks of the previous lactation. The mature equivalent milk production was estimated as equivalent to milk production for a third lactation cow, i.e., 120% of the milk production for primiparous cows and 104% for cows lactating for a second time, as established from the data used by Gaignon et al. (24). Measurement started 3 weeks before the average expected date of calving for each group, on 5 cows of the initial 6, with the extra cow being kept only for blood analyses to replace a cow whose actual calving date might occur too far from the expected date.

The experiment was conducted at the INRA experimental farm of Méjusseaume (longitude −1.71°, latitude +48.11°, Brittany, France) from September 1^st^ 2016 to June 30^th^ 2017. During the experiment, the cows were housed in a free-stall barn cubicle, covered with rubber carpet, except during 3 periods of 3 weeks, for measurements of Ca retention, during which they were transferred to individual tie stalls (1.4 × 2.0 m). The individual stalls were also covered with a rubber mat with individual troughs and individual water bowls. During lactation, a total mixed ration (**TMR**) was distributed twice a day, in 2 equal-sized meals, at 0830 h and 1630 h. The TMR was distributed by an automatic dispenser into the individual troughs, specific to each animal, in the free-stall barn and directly by animal technicians in the individual tie stalls. Cows always had free access to the trough and to water during the day. Lactating cows were fed *ad libitum* and offered quantities calculated to allow 10% orts. Orts were weighed daily before the morning feeding. Cows were milked twice a day, at 0630 h and 1630 h. Milk production and dry matter (**DM**) intake were recorded daily for each individual. Milk composition (fat and protein contents) and somatic cell count were measured twice a week, at the evening and morning milkings. Before calving, each cow was fed a fixed amount of diet, with one distribution per day. Procedures related to care and use of animals for the experiment were approved by an animal care committee of the French Ministry of Agriculture, in accordance with French regulations (project number, 7096-20 16082515505689v2).

### Diets

During the experiment, cows were fed 4 or 5 successive diets according to their physiological stage and the treatment to which they were assigned (Table 1). During the dry period, the offered diets were formulated to cover the requirements for net energy of lactation, protein digestible in the intestine and minerals and vitamins of cows according to the INRA recommendations (19), with restricted quantities offered. The diets offered during the far-off period, i.e., more than 3 weeks before the expected calving date, and the close-up period, i.e., less than 3 weeks before the expected calving date, differed with the specific objective to lower the dietary Ca and the DCAD. For the first 5 d of lactation, all cows received the TMR corresponding to the NCa treatment, whose composition was formulated to cover requirements for net energy of lactation, protein digestible in the intestine, macrominerals, trace minerals, and vitamins of cows according the INRA recommendations (19), with a target DCAD of 200 mEq/kg DM. DCAD is defined as the sum of the diet content of Na^+^ and K^+^ minus the sum of the dietary content Cl^−^ and S^2−^ content, which are expressed in mEq/kg of DM. Five d after calving and until the end of the 10^th^ week of lactation, cows were assigned to one of 3 TMRs corresponding to the compared treatments. The LCa and LCaLD TMRs were formulated to meet the requirements for NE_L_, PDI, and all macrominerals and trace minerals, except for Ca, of cows, according to the INRA recommendations (19). The treatment diets were isocaloric and isonitrogenous with a similar content of P. Differences in the Ca content and DCAD between the 3 TMRs were achieved only by formulating different mineral supplements, with all cows receiving the same base ration.

**Table 1.** Diet centesimal composition and nutritional value.

|  | From -5<br>to<br>-3 weeks | From -3<br>weeks to<br>calving | From 5 to 70 days of lactation |  |  | After 70<br>days of<br>lactation |
| --- | --- | --- | --- | --- | --- | --- |
|  |  |  | NCa | LCa | LCaLD |  |
| Offered amount<br>(kg DM/d) | 12 | 15 | TMR *ad libitum* | TMR *ad libitum* | TMR *ad libitum* | TMR *ad libitum* |
| Ingredients, % of DM |  |  |  |  |  |  |
| **Corn Silage** | 53.7 | 80.5 | 70.0 | 70.2 | 70.1 | 72.1 |
| **Hay** | 14.6 | 0.0 | 0.0 | 0.0 | 0.0 | 0.0 |
| **Wilted and wrapped**<br>**hay (baleage)** | 24.5 | 0.0 | 0.0 | 0.0 | 0.0 | 0.0 |
| **Energy concentrate** | 0.0 | 4.1 | 15.6 | 15.7 | 15.1 | 11.1 |
| **Soybean meal**<br>**(formaldehyde-treated)** | 0.0 | 0.0 | 10.3 | 10.3 | 10.2 | 0.0 |
| **Soybean meal** | 0.0 | 10.1 | 0.0 | 0.0 | 0.0 | 13.1 |
| **Rapeseed meal** | 5.9 | 0.0 | 0.0 | 0.0 | 0.0 | 0.0 |
| **Straw** | 0.0 | 3.3 | 0.0 | 0.0 | 0.0 | 0.0 |
| **Urea** | 0.0 | 0.0 | 1.3 | 1.3 | 1.3 | 0.7 |
| **Mineral Feed** | 0 | 0.7 | 2.9 | 2.3 | 3.3 | 2.5 |
| **Commercial Mineral**<br>**Feed** | 1.3<sup>1</sup> | 1.3<sup>2</sup> |  |  |  |  |
| Mineral Feed g/kg of DM |  |  |  |  |  |  |
| **Calcium Carbonate** | 0.0 | 0.0 | 10.7 | 4.7 | 4.4 | 9.9 |
| **Dicalcium phosphate** | 0.0 | 0.0 | 5.3 | 4.6 | 4.4 | 4.7 |
| **Sodium Sulfate** | 0.0 | 0.0 | 2.7 | 2.3 | 2.3 | 2.4 |
| **Sodium bicarbonate** | 0.0 | 0.0 | 2.2 | 2.8 | 0.0 | 2.4 |
| **Sodium carbonate** | 0.0 | 0.0 | 3.1 | 3.7 | 0.0 | 0.0 |
| **Magnesium oxide** | 0.0 | 0.0 | 0.9 | 0.9 | 0.0 | 1.4 |
| **Hexahydrate**<br>**Magnesium chloride** | 0.0 | 6.7 | 3.6 | 3.7 | 21.5 | 3.8 |
| **Premix<sup>3</sup>** | 0.0 | 0.0 | 0.2 | 0.2 | 0.2 | 0.2 |
| Nutrient Content |  |  |  |  |  |  |
| **CP (g/100 g DM)** | 11.1 | 10.5 | 15.5 | 15.5 | 15.5 | 12.7 |
| **NE<sub>L</sub> (MJ/kg)** | 3.9 | 6.4 | 6.7 | 6.7 | 6.7 | 6.6 |
| **PDI/NE<sub>L</sub> (g/MJ)** | 9.5 | 11.0 | 15.8 | 15.8 | 15.8 | 11.9 |
| **%Ca (g/ kg DM)** | 0.74 | 0.42 | 0.83 | 0.60 | 0.58 | 0.78 |
| **%P (g/ kg DM)** | 0.37 | 0.30 | 0.41 | 0.39 | 0.39 | 0.40 |
| **DCAD (mEq/kg DM)** | 156.8 | 34.8 | 218.0 | 279.5 | 0.0 | 219.9 |

<sup>1</sup> Kéomine Repro (Cooperl Hunaudaye, Lamballe, France): 55.7% Calcium Carbonate, 18.4%
Monocalcium phosphate, 10.0% Magnesium Phosphate, 9% Cane molasses, 2.4 Magnesium
Oxide, 4.5% for trace elements and vitamins.
<sup>2</sup> Kéomine Prépa (Cooperl Hunaudaye, Lamballe, France)
<sup>3</sup> for trace elements and vitamins

### Blood and Milk Sampling

For each group, blood was sampled 3 weeks before the average expected date of calving in the group and at 1, 3, 8, 12, 17, 22, 27 and 31 weeks of lactation (average stage of lactation of the group). After being milked, before being fed, and while restrained in self-locking head gates at the feedline, the cows were sampled for blood by venipuncture of the tail vessels. The samples were collected in vacutainer tubes (Monovette, Sarstedt, Nümbrecht, Germany) coated with lithium heparin for Ca and inorganic P (**Pi**) analyses, and in tubes coated with EDTA for osteocalcin (**OC**) and carboxy-terminal crosslinking telopeptide of type I collagen (**CTX**). OC and CTX are biomarkers of bone formation and resorption, respectively (26). Plasma was recovered after centrifugation at 3,000 *x* g for 12 min within 30 min of sampling and stored at −80°C for OC analysis and at −20°C for other analyses. Milk samples were collected during the morning milking from total milk preceding blood sampling. They were stored at 4°C for analyses of fat and protein contents and for separation of the N, crude protein, soluble and non-soluble Ca and P fractions (i.e., non-protein nitrogen (**NPN**), non-casein nitrogen (**NCN**), urea, soluble Ca and P) and frozen at −20°C for analysis of the total Ca and P contents. Additional samples of milk were also collected at the morning and evening milkings, twice per week for determination of milk fat and protein contents (stored at 4°C before analyses), and on weeks 1, 3, 6, 8, 10, 12, 14, 17, 19, 22, 24, 27, 29 and 31 of lactation, samples were collected for analyses of the milk total Ca and P content (frozen at −20°C before analyses).

### Measurement of Ca and P Retention in Cows

For each group, all input (feed and water intake) and output (excretion in milk, urine and feces) flows of Ca and P were measured 3 times during the experiment, i.e., 3 weeks before the average expected calving date of the group and 3 and 17 weeks after the average calving date. For each measurement, cows were moved from the free-stall barn 2 weeks before beginning the measurements and sent to individual tie stalls for habituation. The individual tie stalls were located in the same building as the free-stall barn. All cows were kept in a same room and were able to smell and hear each other. The feeding modalities remained similar to those applied in the free stall barn. To determine the fecal excretion of Ca and P, large trays were positioned behind the cows on d 15 after the entry of the cows into the stall at 0900 h. Gross fecal output was weighed and sampled from d 16 at 0900 h to d 19 at 0900 h. Two representative samples (500 g fresh each) were dried in a forced air oven (80°C, 72 h) to determine the daily amount of fecal DM excreted. These dried samples were pooled by cow and period for the Ca and P determination. The daily volume of excreted urine was measured from d 15 at 0900 h to d 19 at 0900 h, by equipping cows with urinary catheters connected by a Tygon tube to a 25-L container, which was closed with a rubber plug. To prevent urine deterioration, 250 mL of sulfuric acid (20% vol/vol) was added to the container. The urine was weighed and emptied daily at 0900 h. Each day and for each cow, a sample of 1% of the daily excreted volume was stored at −20°C. At the end of the experiment, these samples were pooled by animal and by period for further Ca and P content analyses.

### Chemical Analysis

Samples of the offered diets, refusals, and feces were ground with a 3-blade knife mill through a 0.8-mm screen. Ash was determined by calcination at 550°C for 5 h in a muffle furnace. Nitrogen concentration was determined by the Dumas method, according to the Association Française de Normalisation (AFNOR, 1997), on a LECO apparatus (LECO, St. Joseph, MI). The dietary, fecal, urine and milk Ca contents were measured by atomic absorption spectrophotometry (Spectra-AA20 Varian, Les Ulis, France), with use of lanthanum chloride solution to dilute the sample and after calcination of the solid samples (500°C for 12 h). Phosphorus contents were determined using a KONE PRO multi-parameter analyzer (Thermo Fisher Scientific, Illkirch, France) by the Allen method for P (27). Milk fat and protein contents were determined by a commercial laboratory using mid-infrared analysis (Mylab, Châteaugiron, France). Milk content of total N (Kjeldahl), nonprotein N (precipitation at pH 4.6 with trichloroacetic acid and filtration), NCN (precipitation at pH 4.6 with 10% acetic acid and 1 M sodium acetate) content, and urea (colorimetric analysis) were determined according to the methods described in Hurtaud et al. (28). Plasma OC and CTX concentrations were determined by ELISA with a CrossLaps kit from IDS (Paris, France) for CTX and a kit from Quidel (San Diego, CA) for OC (Inter-assay variation: 3.7%, intra-assay variation: 3.3% for CTX: Inter-assay variations: 5.42%, intra-assay: 7.35 for OC).

### Statistical Analysis

Data were analyzed using PROC GLIMMIX in SAS (29), with a generalized linear mixed model with repeated values given here:

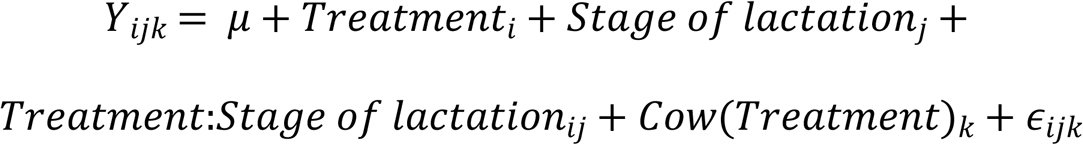

where *Y_ijk_* was a dependent variable of a cow *k*, within treatment *i* at the stage of lactation *j*. Treatment, stage of lactation and their interaction were fixed factors, and the cow was a random factor. Measurements repeated over the stage of lactation were considered by using a covariance matrix. The choice of the structure of the matrix was determined according to the structure of the data and then was performed using an AIC for analyses of all variables. Average flows of Ca and P during the 4 d of measurements were also analyzed independently during each stage of lactation with a generalized linear model using proc GLM in SAS and included only the fixed effect of the treatment. These analyses supplemented the previous analyses, performed with the complete model because it has been shown that with a lower number of data, the inclusion of repeated measurements in a generalized linear mixed model can deteriorate the quality of detection of significant effect by increasing the second species risk (30).

### Results

The distribution of the cows among the 3 treatments had to be modified before the differentiation of the TMR 5 days after calving. Two cows calved more than 2 weeks before the expected calving: one was assigned to the NCa treatment; the other, to the LCaLD treatment. They were removed from the experiment and replaced by the extra cows, kept in each of the considered group, and only one blood sampling before calving was performed for each replacement cow. Another cow, initially assigned to the LCaLD treatment and diagnosed with a milk fever, was replaced by a cow initially assigned to the NCa treatment and belonging to the same group of calving date. At 7^th^ week of lactation, one cow of the NCa treatment died due to bowel obstruction, and its data were removed from the data set. Despite these events, the pre-experimental characteristics of the 3 experimental lots remained similar. Average parities were 2.4 for the LCaLD and 3.0 for the TEM and LCa. Milk production during the first 32 weeks of the previous lactation was not affected by the treatments (Fig 1A, P = 0.92 for the treatment effect, and P = 0.93 for the effect of the interaction treatment × stage of lactation). Mature milk yield was not different between the treatment either (Fig 1B, P = 0.99 for the treatment effect, and P = 0.95 for the effect of the interaction treatment × stage of lactation). The average mature equivalent milk productions over the first 32 weeks of lactation were 33.6 (± 3.94), 33.2 (± 3.52) and 33.1 (± 3.52) kg/d for the treatments NCa, LCa and LCaLD, respectively. Neither milk protein content (Fig 1C, P = 0.62) nor milk fat content (Fig 1D, P = 0.58) were affected by the treatments during the first 32 weeks of the previous lactation, but individual variability of those last parameters was high. The average milk protein contents over the 32 first weeks of lactation were 29.8 (± 0.97), 31.0 (± 0.87) and 31.0 (± 0.87) g/kg for the treatments NCa, LCa and LCaLD, respectively. The average milk fat contents over the first 32 weeks of lactation were 36.8 (± 2.48), 40.3 (± 2.22) and 39.3 (± 2.22) g/kg for the treatments NCa, LCa and LCaLD, respectively.

**Fig 1.**
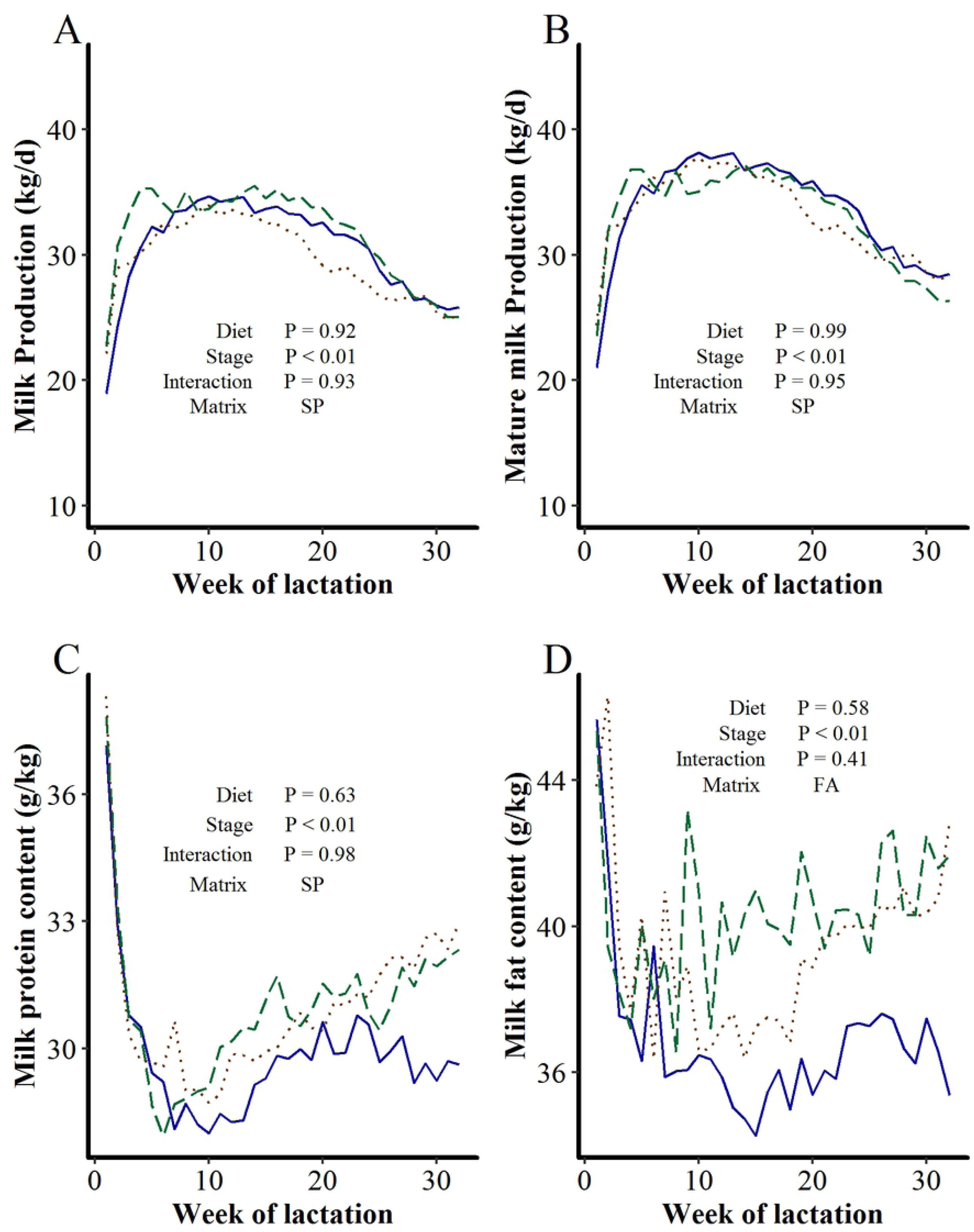
Differences between cows according to their treatments on A) milk production, B) mature equivalent milk production, C) milk protein content and D) milk fat content during the first 32 weeks of the lactation preceding that described in the present paper. NCa: normal line, LCa: dashed line, and LCaLD: dotted line

### Ca, P and DM intake

During the period that the experimental diets were fed, from 5^th^ to 70^th^ day of lactation, the Ca intake was significantly lower with the LCa and LCaLD treatments compared with NCa (P < 0.01). During this period, the average daily intake of Ca was 136.0 (± 4.98) g/d and 126.5 (± 4.98) g/d for LCa and LCaLD, respectively, and 184.1 (± 5.06) g/d for NCa, leading to Ca intake that was 31% lower for the LCa and LCaLD treatment compared with the NCa, as expected. During the same period, the difference between absorbable Ca intake and Ca requirements calculated according the INRA feeding system (19), was negative for the LCa and LCaLD treatments (−13.4 ± 1.36 and −15.5 ± 1.36 g/d, respectively), whereas it was positive for the NCa treatment (5.4 ± 1.42 g/d, P < 0.01, Fig 2A). After 70^th^ day of lactation, when cows received the same ration, the Ca intake was not affected by the treatments, and the difference between absorbable Ca intake and Ca requirements always remained positive, with an average value of 10.9 ± 1.12 g/d (Fig 2A). The DM intake tended (P = 0.08, Fig 2B) to be higher for LCa compared with the LCaLD and NCa treatments, with DM intakes of 23.0 ± 0.4, 21.6 ± 0.4 and 22.0 ± 0.5 kg/d for the treatments LCa, LCaLD and NCa, respectively, throughout the lactation. Consequently, the P intake also tended (P = 0.08, see Table S1) to be higher with LCa compared with the LCaLD and NCa treatments across lactation, with P intake of 91.8 ± 1.70, 86.1 ± 1.7 and 89.4 ± 1.90 g/d for the treatments LCa, LCaLD and NCa, respectively. The amount of P ingested by the cows exceeded the requirement by 4.09 ± 0.90 g/d during the first 70 days of lactation.

**Fig 2.**
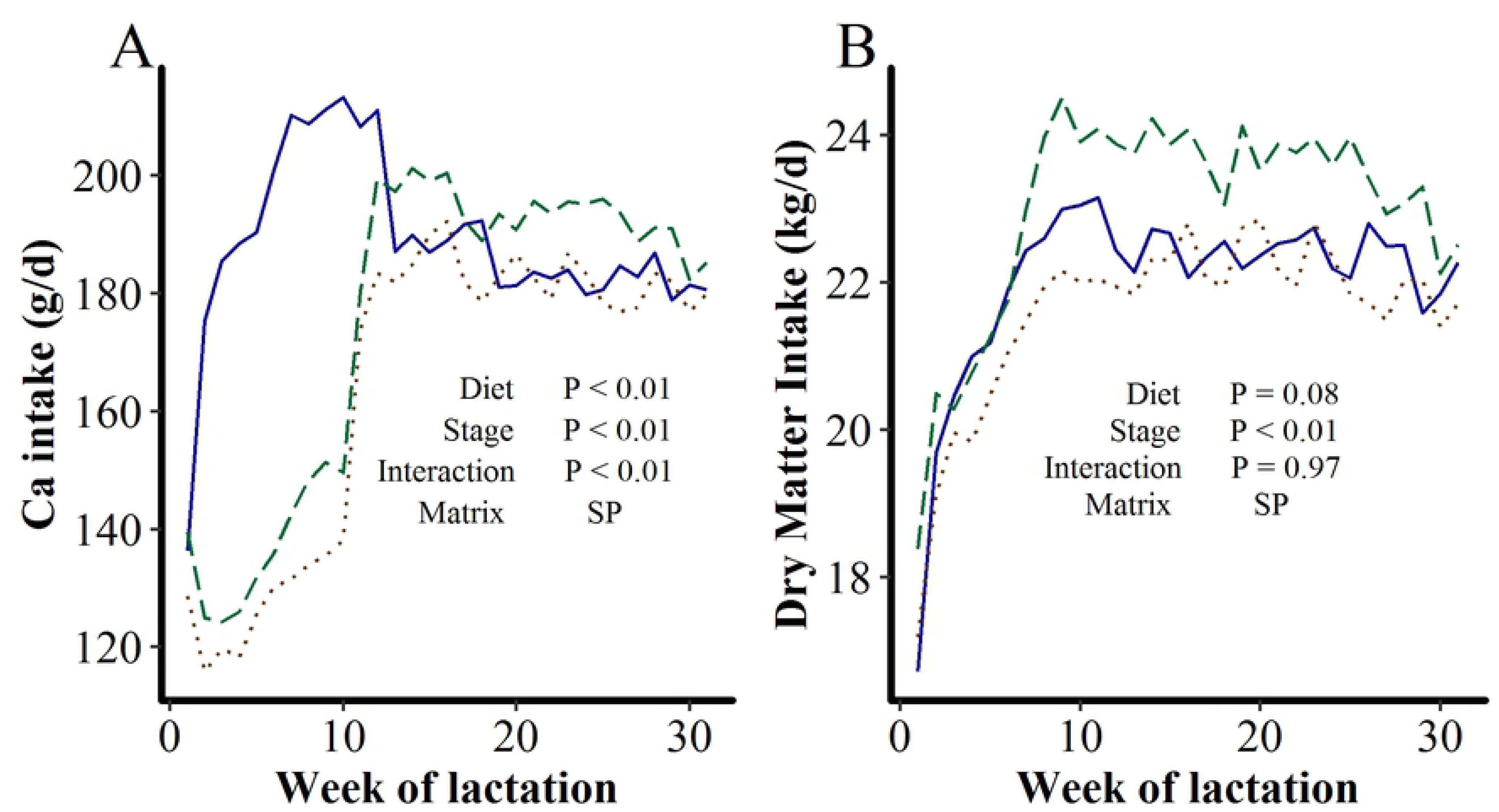
Effect of dietary content of Ca and DCAD between days 5 and 70 of lactation on A) Difference between the estimated supply and requirement of absorbable Ca and B) Dry matter intake. NCa: normal line, LCa: dashed line, and LCaLD: dotted line

### Plasma Concentrations of Biomarkers of Bone Formation and Resorption, Ca and Pi

The plasma concentrations of OC and CTX were affected neither by the treatments nor by the interaction treatment × stage of lactation (Fig 3). For all treatments, plasma OC concentrations decreased after calving to reach a minimal value one week after calving, whereupon it increased sharply for 10 weeks and then more slowly until the end of the lactation (stage of lactation, effect P < 0.01). For all treatments, the plasma CTX concentrations increased sharply after calving to reach a maximal value between week 3 and 8 of lactation and then decreased until the end of lactation (stage of lactation effect P < 0.01). The plasma Ca concentration was, on average, 100.3 (± 1.84) mg/L, and individual values always remained between 80 to 120 mg/L, with only one cow in hypocalcemia at one week of lactation (76 mg/L). Plasma Ca concentration tended to be lower at 1 and 3 weeks of lactation compared with the other sampling times (stage of lactation effect P = 0.06, see Table S1) but was affected neither by the treatments (P = 0.63) nor by the interaction treatment × stage of lactation (P = 0.31). Plasma Pi concentration was 50.7 (± 1.78) mg/L on average. Some individual values could be lower than 40 mg/L at 3 weeks of lactation, but no individual data were higher than 80 mg/L. Plasma Pi sharply decreased after calving, at 1 week and 3 weeks of lactation, increasing afterward and leveling off at 17 weeks of lactation (stage of lactation effect, P < 0.01, see Table 1). It was affected neither by the treatment (P = 0.89) nor by the interaction treatment × stage of lactation (P = 0.63).

**Fig 3.**
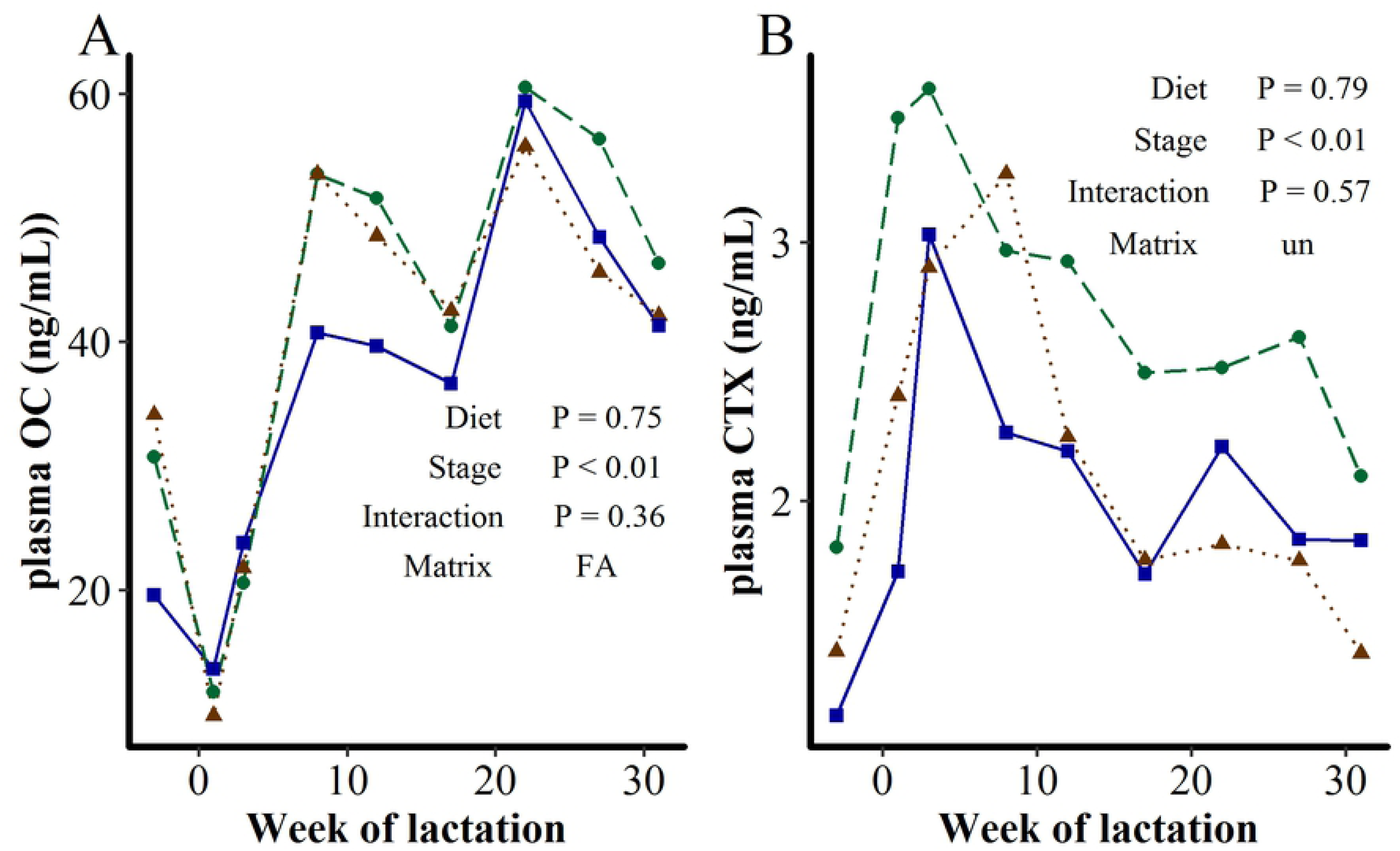
Effect of dietary content of Ca and DCAD between d 5 and d 70 of lactation on A) plasma OC concentration and B) plasma CTX concentration. NCa: normal line, LCa: dashed line, and LCaLD: dotted line

### Ca and P Partitioning and Retention

During the 4 d of measurement of Ca and P retention, the Ca intake was lower for the LCa and LCaLD treatments compared with the NCa at 3 weeks of lactation, i.e., during the period of TMR differentiation between treatments (P < 0.05, Fig 4A), whereas the Ca intake was not affected by the treatment 3 weeks before calving and at 17 weeks of lactation (P > 0.10 for both stages, interaction treatment × stage of lactation P < 0.001). The daily amount of Ca excreted in the feces was also lower for the LCa and LCaLD treatments compared with the NCa at 3 weeks of lactation (P < 0.001, Fig 4B) and was not affected by the treatment 3 weeks before calving and at 17 weeks of lactation (P > 0.10 for both stages, interaction treatment × stage of lactation P < 0.001). Apparent digestibility of the Ca increased after calving (Fig 4C, P < 0.001) from 21.0% (± 2.33) 3 weeks before calving to 36.4% (± 2.10) and 33.4% (± 2.10) at 3 and 17 weeks of lactation, respectively. Apparent digestibility of Ca was not affected by the treatments 3 weeks before calving and at 17 weeks of lactation but it was higher for the LCa and LCaLD treatments compared with NCa at 3 weeks of lactation (P = 0.03, average of 37.3 ± 3.5% for LCa, 41.7 ± 3.5% for LCaLD and 30.1 ± 3.9% for NCa). Daily amounts of Ca excreted in urine were low, 2.00 g/d on average, compared to the other flows of the input-output retention, as expected, and this flow tended to be affected only by the stage of lactation (P = 0.09, Fig 4D). However, the daily amount of Ca excreted in the urine was higher with the LCaLD treatment compared with the NCa treatment and the LCa treatment at 3 weeks of lactation (P < 0.05, interaction treatment × stage of lactation P = 0.004, 4.4 ± 0.5 g/d for LCaLD vs. 0.5 ± 0.6 g/d for NCa and 0.7 ± 0.5 g/d for LCa). The daily amount of Ca secreted in milk was on average 48.2 ±1.42 g/d at 3 weeks of lactation and 42.7 ±1.42 g/d at 17 weeks (Fig 4E). The amount of Ca secreted in milk daily decreased slightly between 3 and 17 weeks of lactation (P < 0.001) and was not affected by the treatments at any stage. Daily Ca retention, i.e., the difference between input and output of Ca, was very positive before calving (+17.3 ± 2.84 g/d, Fig 4F), decreased at 3 weeks of lactation to an average value close to 0 (2.1 ± 2.62 g/d) and increased at 17 weeks of lactation to a positive value (+16.7 ± 2.62 g/d, stage of lactation P < 0.001). At 3 weeks of lactation, the daily Ca retention tended to be lower for the LCa and LCaLD treatments compared with the NCa treatments (P = 0.09), with values around equilibrium for the LCa and LCaLD (−2.1 ± 4.3 g/d for LCa, + 0.3 ± 4.3 g/d for LCaLD and + 8.1 ± 4.8 g/d for NCa). The daily Ca retention was not different between the 3 groups of cows dedicated to the 3 treatments 3 weeks before calving and at 17 weeks of lactation (P > 0.10).

**Fig 4.**
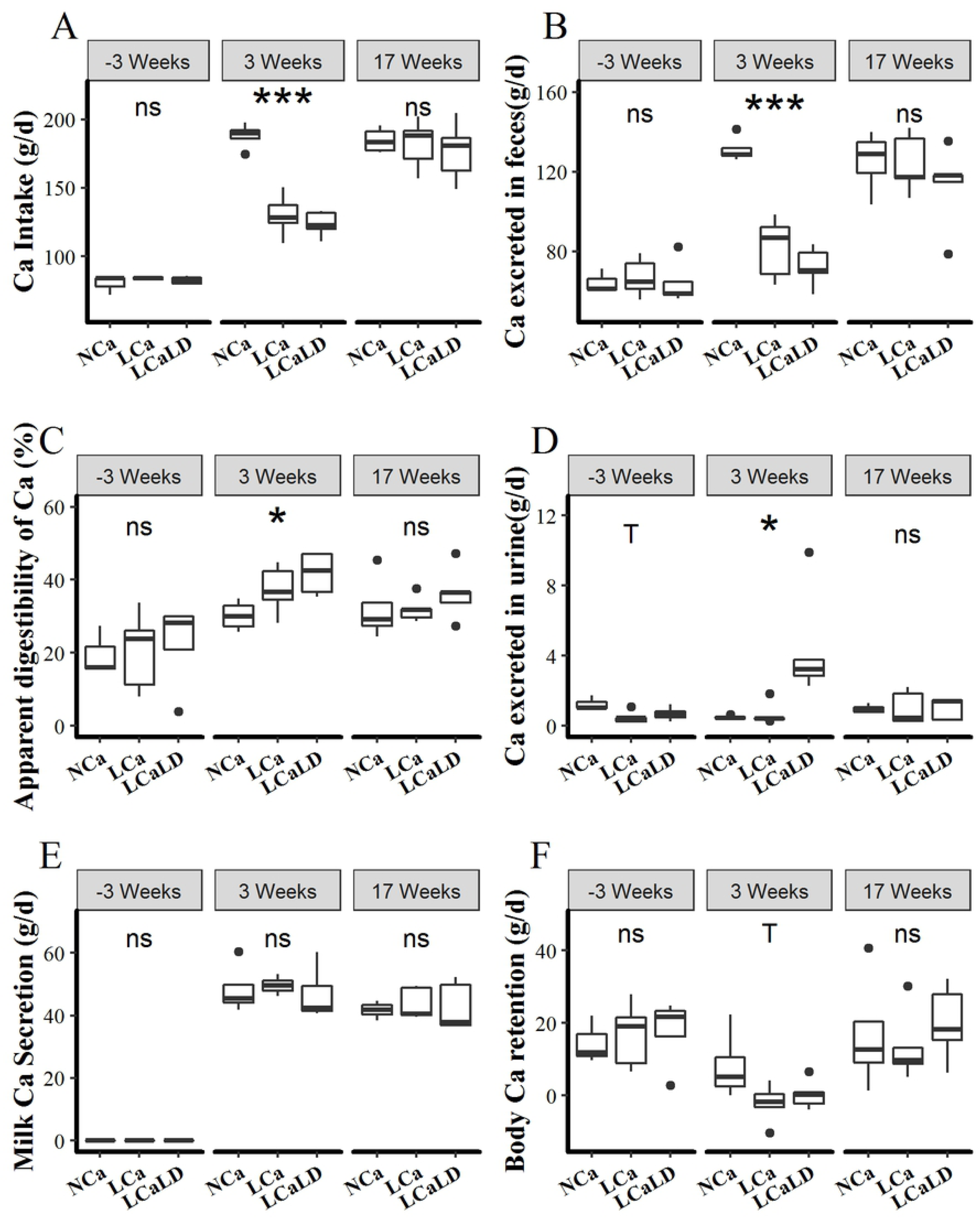
Effect of dietary content of Ca and DCAD between d 5 and d 70 of lactation A) daily Ca intake, B) fecal losses of Ca, C) Ca secretion in milk, D) urinary losses of Ca, E) apparent digestibility of Ca and F) Ca balance. Boxplots are obtained from mean data per animal. Signs give p-values of the dietary effect for intra-period analyses: ns = not-significant (P > 0.10); T = Tendency (P<0.10), * = significant (P<0.05), ** = significant (P < 0.01); *** = significant (P < 0.001)

Neither the DM intake, intake of P, fecal or the excretion of DM and P were affected by the treatment or the interaction treatment × stage of lactation (P > 0.10, see Table 1). Both the apparent digestibility of the DM and P increased at calving and leveled off after 3 weeks of lactation. Apparent DM digestibility was 65.1 ± 0.86 at 3 weeks before calving, 72.7 ± 0.79% at 3 weeks of lactation and 71.6 ± 0.79% at 17 weeks of lactation (P < 0.001), and apparent P digestibility was 10.9 ± 2.17% 3 weeks before calving, 54.7 ± 1.98% at 3 weeks of lactation and 47.5 ± 1.98% at 17 weeks of lactation (P < 0.001). Daily P retention tended to increase at 17 weeks of lactation compared with both other stages of lactation, with values of 4.3 ± 1.66 g/d at 3 weeks before calving, 7.9 ± 1.48 g/d at 3 weeks of lactation and 10.0 ± 1.48 g/d at 17 weeks of lactation (P = 0.08).

### Milk Production and Composition

Milk production tended to be lower throughout the 32 weeks of lactation for treatments LCa and LCaLD compared with treatment NCa (P = 0.09, Fig 5A), with average values of 36.8 ± 0.9 kg/d for LCa, 35.9 ± 0.9 kg/d for LCaLD and 39.2 ± 1.1 kg/d for NCa. This production led to a difference in cumulative milk production at 200 d of lactation between the low Ca treatments (LCa and LCaLD) and the control treatment NCa of more than 400 kg. The difference in milk production between the low Ca treatments and NCa was maximal at the 4^th^ week of lactation, with a milk production difference of more than 4.5 kg/d. Milk Ca content sharply decreased to reach a minimal value at 3 weeks of lactation (Stage of lactation effect, P < 0.01, Fig 5B). Then, after 8 weeks of lactation, the milk Ca content increased for the LCa and LCaLD treatments, whereas it remained stable for NCa (interaction stage of lactation × treatment P < 0.01 in the morning, non-significant in the evening). After the 24^th^ week of lactation, the morning milk Ca contents were 1230 (± 48.4) and 1228 (± 48.4) mg/kg for the LCa and LCaLD treatments, respectively, and 1128 (± 54.2) mg/kg for NCa (P < 0.01). Milk protein content was numerically higher for cows fed with the treatments LCa and LCaLD, compared with those fed the NCa, between 5 and 9 weeks of lactation and after 20 weeks of lactation, but the treatment effect was not significant (Fig 5C). Milk fat content was also unaffected by the interaction of stage of lactation × treatment (Fig 5D, P > 0.10).

**Fig 5.**
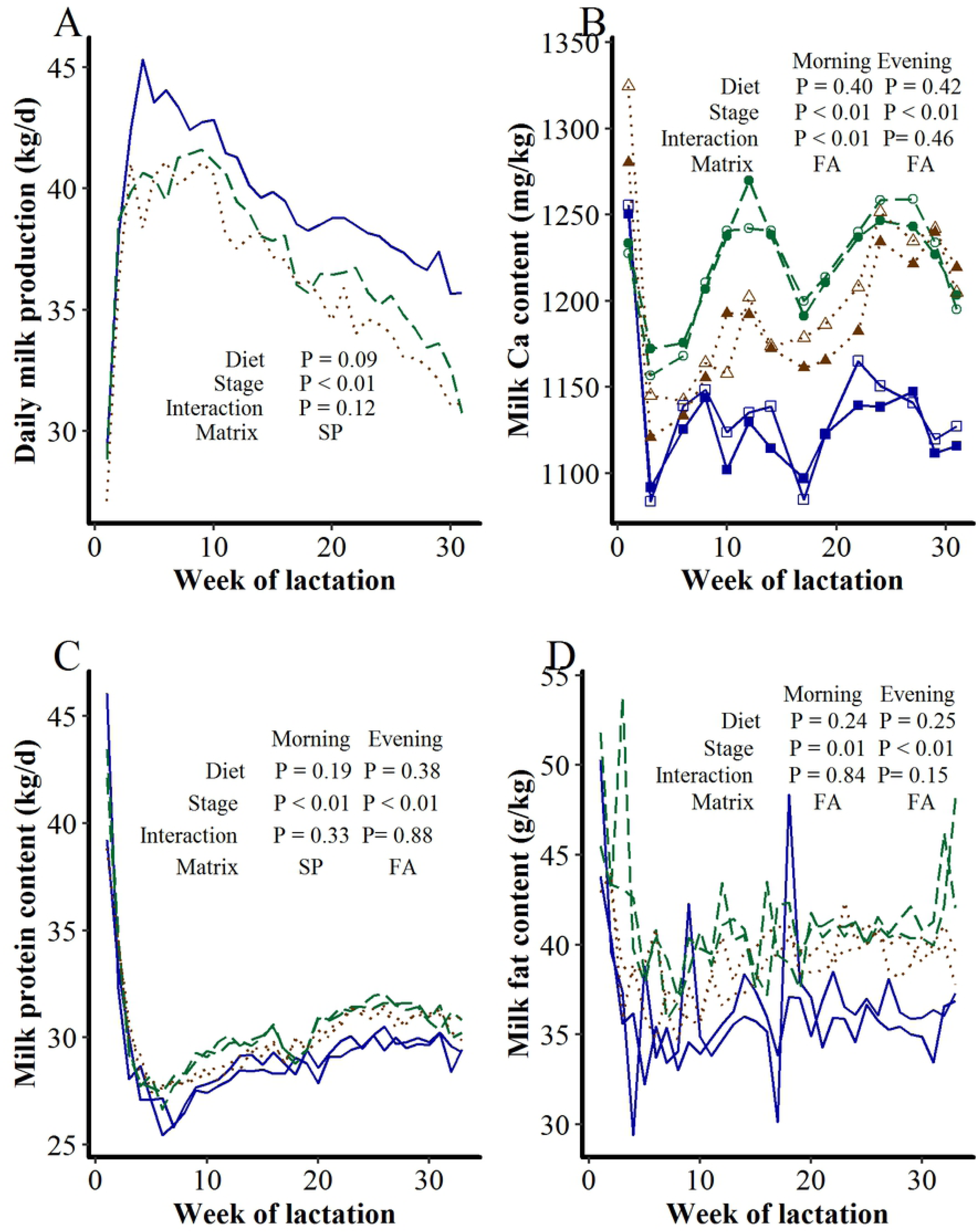
Effect of diet content of Ca and DCAD between d 5 and d 70 of lactation on A) daily milk production, B) milk Ca content, C) milk protein content and D) milk fat content. NCa: normal line, LCa: dashed line, and LCaLD: dotted line. For milk Ca content, color-filled shapes are for morning milk Ca content and the white-filled shapes for evening milk Ca content.

### Milk Ca and Protein Partitioning between Soluble and Colloidal Phases

Milk casein content was higher with the LCa and LCaLD treatments compared with the NCa (Fig 6A, P < 0.001), with this difference being more distinct after 17 weeks of lactation (interaction stage of lactation × treatment, P < 0.01). At the end of the period of diet differentiation between treatments, the milk casein content of the NCa treatment increased transitorily. The ratio between the milk contents of colloidal Ca and casein increased at the beginning of the lactation and remained relatively stable after 8 weeks of lactation at a value approaching 36 mg/g (Fig 6B). This ratio was not affected by either the treatments or the interaction stage of lactation × treatment (P > 0.10). The proportion of soluble Ca among total Ca was lower for the LCa and LCaLD treatments compared with the NCa (Fig 6C, P < 0.01). This proportion decreased transitorily at the end of the period of diet differentiation between treatments for the NCa treatment (Interaction stage of lactation × treatment, P = 0.03). On average, 27% of the milk Ca was in the soluble form. The ratio between the milk Ca and protein contents increased at the beginning of the lactation to peak at approximately 6-8 weeks of lactation and then decreased to level off after 17 weeks of lactation to a value close to 39 mg/g on average (Fig 6D, P < 0.001). It was affected neither by the treatment nor by the interaction stage of lactation × treatment.

**Fig 6.**
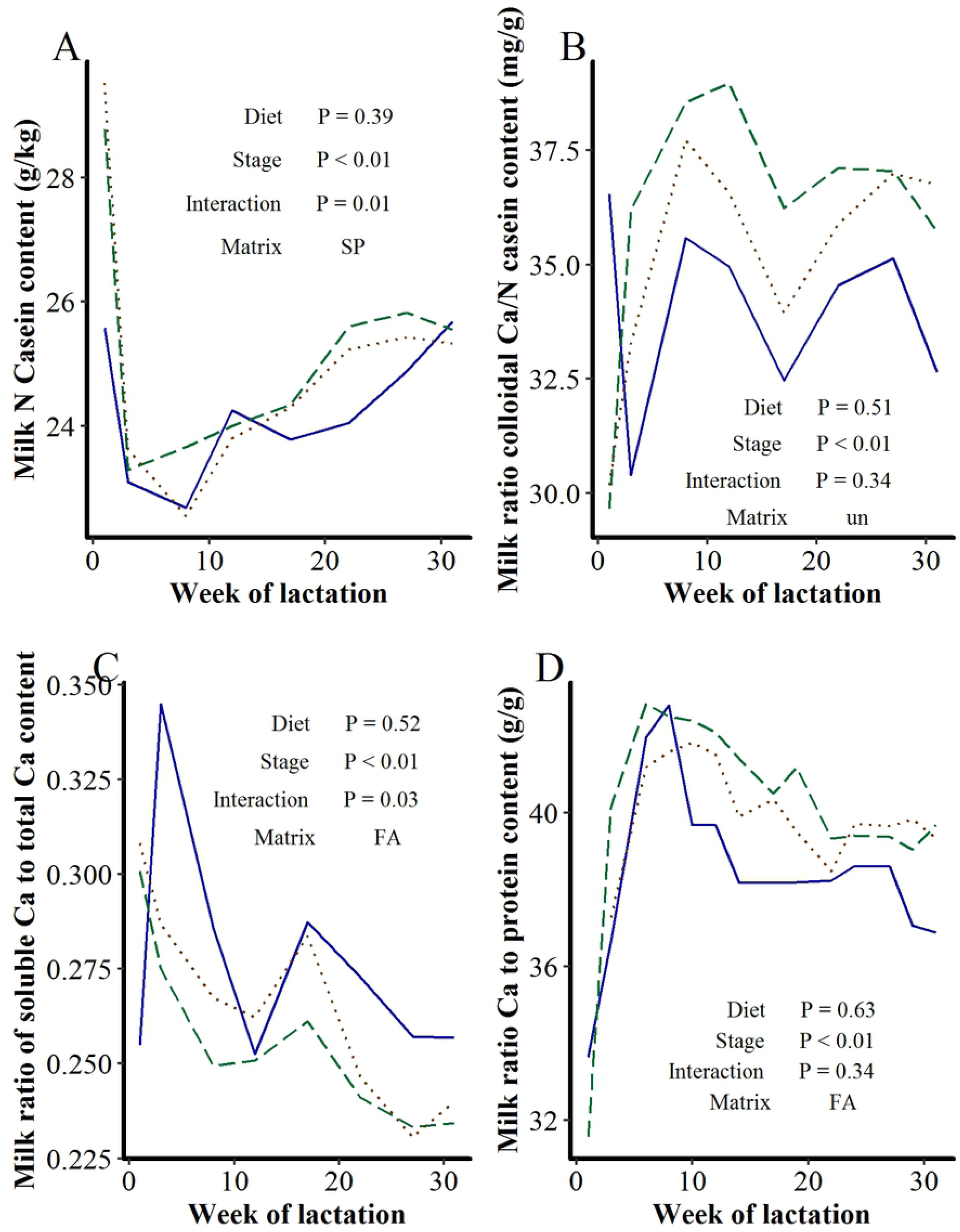
Effect of dietary content of Ca and DCAD between d 5 and d 70 of lactation on morning milk A) Casein content, B) ratio of colloidal Ca to casein content, C) ratio of soluble Ca to total Ca content and D) ratio Ca to protein content. NCa: normal line, LCa: dashed line, and LCaLD: dotted line.

## Discussion

### A limited effect of dietary Ca supply and DCAD on the dynamics of bone mobilization and reconstitution during lactation

The objective of the treatments LCa and LCaLD was to induce an increased bone mobilization during the first 10 weeks of lactation. For this purpose, the dietary Ca supply was limited to 70% of the recommended supply to cover the cows’ requirements according to the expected milk production and intake according to the INRA feeding system (INRA, 2010). With such restriction of Ca supply, some studies highlighted a decrease in the body retention of Ca at the beginning of lactation in dairy cows, with this decrease in body Ca retention later reaching negative values (11,31), whereas some studies highlighted an increase in the serum concentration of pyridinoline, which is a biomarker of bone resorption (16). Both results suggested an increase in mobilization of Ca from bones. To increase the chance to induce a bone mobilization in our experiment, the DCAD was also decreased for treatment LCaLD. The positive effect of low DCAD on bone mobilization is used to prepare the bones of cows to be mobilized before lactation for milk fever prevention (32). An expected effect of lowering the DCAD is to decrease the pH of the blood, to avoid changes in the PTH receptor conformation, and to preserve the receptor affinity for PTH (32). Another expected effect is that bone can release Ca from bone carbonates when the local pH is low as a buffering system (33). The DCAD of the LCaLD treatment in our experiment approached 0, which is a maximal limit under which a positive effect on the prevention of milk fever, and perhaps even on bone mobilization, could be expected (34). In our experiment, the diet offered to the cows during the first 10 weeks of lactation was also intentionally formulated with a high dietary PDI to NE_L_ ratio, with the objective being to maximize the milk production capacity by the mammary gland and perhaps the necessity of bone mobilization (20). Primiparous cows were excluded from the experiment, and special care was applied to balance the average parity between treatments because parity is, in dairy cows, a strong determinant of bone formation and resorption throughout lactation (11,25,35) and digestive absorption of Ca (36).

However, treatments LCa and LCaLD, compared with NCa, only tended to induce a small decrease in Ca retention 3 weeks after calving, with no effect on the dynamics of the blood biomarkers of formation and resorption throughout the lactation. Even at 3 weeks of lactation, the Ca retention remained close to 0 with the treatments LCa and LCaLD and was not clearly negative as it might have been expected from other studies (11,31). A debate exists about the effect of a low supply of Ca and P on the amplitude of bone mobilization at the beginning of lactation. The results of Braithwaite (15) illustrated that a strong restriction of the Ca and P supply did not affect the amount of Ca mobilized from bone at the beginning of lactation of ewes but strongly reduced the amount of Ca retained in bone for bone reconstitution at the end of lactation. Benzie et al. (4) also showed that a restriction of the Ca supply lowered the mineral contents of the bones of ewes slaughtered at 100 d of lactation. These results suggest that bone mobilization at the beginning of lactation was programmed in response to homeorhesis regulation, which was linked to the stage of lactation, and the level of the Ca supply did not affect it. Because we observed only a limited effect of restriction of Ca dietary supply on bone mobilization at the very beginning of lactation, i.e., at 3 weeks of lactation, the possibility exists of such regulation. However, the data of Taylor et al. (11) illustrated, in Holstein dairy cows, a clear effect of dietary Ca supply on the body retention of Ca even at 2 to 5 weeks of lactation. With Ca supplies approaching those of the low Ca treatments in our experiment, these authors measured Ca retention of −14.5 g/d, whereas with Ca supplies approaching those of the NCa treatment, the measured Ca retention was 7.4 g/d. With high Ca supply, i.e., 1.03 g/kg MS, the Ca retention they measured was even as high as 31.7 g/d. These observations contradicted the hypothesis that bone mobilization at the beginning of lactation would be programmed in response to homeorhesis regulation linked to the stage of lactation independently from the level of Ca supply. After 20 weeks of lactation, as observed by Taylor et al. (11) and Braithwaite (15), at the end of lactation in ewes, the Ca supply also affected the bone reconstitution, with a proportionally higher Ca retention with diets providing more Ca. In our experiment, we did not observe any effect of the treatment on the Ca retention at 17 weeks of lactation. This result may be explained by the fact that the dietary supply was not different among the treatments after 10 weeks of lactation, that is during a stage preceding the period of end of bone mobilization, i.e., 3 months in dairy cows (2). This might suggest that the immediate Ca supply may be more important than the bone status to drive the amplitude of the reconstitution. However, the difference in the Ca mobilization among the treatments in our experiment may have been too low to significantly affect the status of bone reserves at 17 weeks of lactation.

The dynamics of the blood bone biomarkers throughout lactation observed in our experiment agreed with those previously observed (10,11,13,20), with a sharp decrease in OC being observed at calving and an increase in CTX being observed at the beginning of lactation. These results agreed with the measurements showing lower Ca retention at the beginning of lactation compared to that measure before calving or at 17 weeks of lactation, illustrating a net bone reconstitution at those times.

### Dairy cows have adapted to low dietary Ca supplies by increasing digestive absorption of Ca in early lactation

In our experiment, as with that of Taylor et al. (11) and Braithwaite (15), the evolution of calcemia during the lactation-gestation cycle was not affected by the dietary Ca supply, which confirms that calcemia is very tightly regulated. This suggests that, if bone mobilization was not the main effector mobilized for the regulation of calcemia when the Ca supply was lowered in our experiment, other Ca flows must have allowed this regulation. Our results clearly illustrate that the decrease in Ca intake with the treatments LCa and LCaLD was almost entirely compensated, at the scale of the organism, by an equivalent decrease in the daily amount of Ca excreted in feces, i.e., by an increased apparent digestive absorption of Ca. The apparent digestibility of Ca was even quite high, with an average higher than 40% for the LCaLD. These results contrasted with those of Taylor et al. (11) and Moreira et al. (16), who observed lower apparent digestibility of Ca at a similar stage of lactation, with highest values at approximately 35%. A reason for that may be that a significant proportion of the dietary Ca was provided by alfalfa silage or hay in those studies, whereas it was mainly provided by a mineral source of Ca in ours. Ca from alfalfa is known to be less available for absorption in ruminants (5). Possibly, in contrast to our findings, these authors could observe an increase in bone mobilization under conditions of low calcium supply at approximately 3 weeks of lactation because the cows could not increase their apparent absorption of Ca because of the low availability of dietary Ca in their feed. In our experiment, the cows may have experienced an increase in digestive absorption rather than a mobilization of Ca from bone to regulate calcemia because dietary Ca was more available for absorption. This hypothesis would require confirmation. The effect of the dietary Ca content on the apparent or real absorption of Ca had been clearly illustrated by (37) on nonpregnant, nonlactating cows on diets that did not contain alfalfa. A clear increase in bone mobilization at the beginning of lactation with low dietary supply of Ca has also been observed by (15) in ewes. However, these authors also observed a concomitant increase in the digestive absorption of Ca. The reason for the adaptive mechanisms to co-exist in this study in contrast to our study may be explained by the very important restriction of Ca supplied compared to that in the experiment of Taylor et al. (11), Moreira et al. (16) or ours. A high restriction of Ca likely necessitates the implementation of more adaptive mechanisms.

We clearly observed a strong effect of the physiological stage of the cows on the apparent digestibility of Ca, with higher digestibility at 3 or 17 weeks of lactation compared with 3 weeks before calving. The increase in the absorption capacity of Ca by the digestive tract between gestation and lactation agrees with the increase in PTH release and 1,25-(OH)_2_D_3_ synthesis at the onset of lactation (2) but has not been so clearly illustrated. We also observed a clear positive effect of the LCaLD treatment on the daily amount of Ca excreted in urine, but the size of the flow, i.e., less than 4 g/d at 3 weeks of lactation, is limited compared with that of Ca excreted in milk, i.e., approximately 50 g/d or in feces, approximately 100 g/d. The increase in this flow with the treatment providing a DCAD close to 0 agrees with the idea that high hydrogen ion content in the glomerular filtrate interferes with the ability of the kidneys to reabsorb Ca from the filtrate, causing the increase in urinary Ca excretion (32).

### The relation between the dynamics of milk Ca content and bone formation and resorption during lactation

Experiments with lactating mice have demonstrated that a decrease in Ca intake can induce an increase in bone resorption and a concomitant decrease in the milk Ca to protein ratio (22). Those effects have been shown to be mediated by the CaSR of the epithelial cells of the mammary gland, with a lack of Ca on the CaSR decreasing Ca transport into the milk and increasing PTHrP secretion by the mammary gland, and thus bone resorption. Our hypothesis was that low Ca intake would induce both an increase in bone resorption and a decrease in the secretion of Ca in milk. However, the effects of the low Ca supply in our experiment had only a limited effect of bone mobilization at the beginning of lactation; thus, this experiment did not fully allow testing our hypothesis.

The cows affected by the LCa and LCaLD treatments tended to have higher milk Ca content after 10 weeks, which did not agree with our hypothesis for 2 reasons. First, this difference appeared when the diet no longer differed by treatment. Second, according to our hypothesis, a lower milk Ca content was expected for those treatments. Because the genetics of the cow is known to be a major determinant of milk Ca content in cows (38) and because the milk Ca content was not measured prior to the previous lactation when attributing cows to the treatments, possibly the cows of the NCa treatments had lower milk Ca contents because of their genetics. The milk casein contents of these cows was also lower, and most milk Ca is bound to casein (39), with a stable ratio between colloidal Ca and casein, as observed in our experiment. However, we could also observe a very transient change in the proportion of the milk-soluble Ca and even of the total milk Ca contents when the TMR changed at 10 weeks of lactation milk. This result suggests that milk Ca content may be related to the Ca homeostasis of cows in a very transient way on the first day of perturbation. Detecting and fully explaining such quick change with a sampling interval of 2 weeks is impossible. This would indicate that the milk Ca content may not be an indicator of the whole shape of bone resorption at the scale of the lactation.

### A possible effect of restricted Ca intake on milk production and cow longevity?

The difference of 2 kg/d in milk production between the NCa treatment on one side and the LCa and LCaLD treatments on the other side was an unexpected result that could not be attributed to the measured pre-experimental characteristics of the cows. The discrepancy between the treatments appeared approximately 2 weeks after the differentiation of the diets according to the treatment, i.e., at 3 weeks of lactation, and lasted until the end of the experiment, i.e., largely after 10 weeks of lactation, when all cows began to receive the same diet. It is impossible to draw firm conclusions about the effect of dietary Ca supply on milk production from our experiment given the low number of cows involved. However, it cannot be totally excluded either that, the low Ca intake did not impair the potential milk secretion by the mammary gland at peak lactation by altering either the proliferation of the mammary epithelial cells or their exfoliation. Ca has been shown to be involved in cell proliferation as an important messenger, notably for the breakdown of the nuclear envelope (40). Possibly, the milk production potential of cows in the LCa and LCaLD was not totally expressed because Ca may have limited the capacity for cell proliferation during early lactation. Wohlt et al. (31) also observed a decrease in the milk production of cows with lower Ca supply, with dietary Ca contents between 0.9 and 0.6% DM, but the response depended on the Ca source. Taylor et al. (11) did not observe any effect of the dietary Ca contents on milk production but two-thirds of the cows were primiparous. Moreira et al. (16) did not observe such an effect either, with multiparous cows, but their experiment stopped after one month of lactation. Older studies (41) highlighted an effect of Ca supplementation on milk production of dairy cows, but bone meal was used as Ca supplementation, and bone meal also contained P, which is known to affect the DM intake and milk production. Possibly, because we used a high level of protein supplementation, with formaldehyde-treated soybean meal partially protected from protein ruminal degradation, and multiparous cows, the milk production potential may have been maximized and the Ca could have been a limiting factor, but this remains to be confirmed on more animals.

Another unexpected observation in our experiment was that the culling rate of cows before the next calving was numerically clearly higher in the LCa and LCaLD treatments compared with the NCa treatments. With the LCa treatment, 3 of the 5 cows were culled before the next lactation, 1 because of the absence of estrus detection, and 2 because of claw disorders. With the LCaLD treatment, 1 of the 5 cows was culled because of failures to be artificially inseminated. All cows from the NCa treatment were kept for subsequent lactation without health or reproductive problems before the calving. Due to the low number of cows involved in this experiment, affirmation of an effect of the dietary Ca content on cow’s reproduction and health from our results was not possible. The subclinical hypocalcemia during the first three days of lactation has a negative effect on the reproductive performances of cows. Our results suggest that Ca supply under the requirements during the first weeks of lactation could also have a detrimental effect (42), but this remains to be demonstrated with more animals.

## Conclusion

Lowering the dietary Ca content to between 0.8 and 0.6 g/kg dry matter, clearly increased the apparent digestive absorption of Ca of the cows at 3 weeks of lactation but marginally affected the body retention that remained nearly zero. This result suggests that bone mobilization of cow at the beginning of lactation can be unaffected by the supply of Ca, as long as the source of Ca is available for absorption. The low supply of Ca did not clearly affect the milk Ca content. However, it cannot be excluded that it did not lower the milk production and did not affect the reproductive performances of the cows and their probability to continue for a subsequent lactation. These results need to be confirmed using a higher number of animals, while also suggesting that Ca supplementation must be carefully checked at the beginning of lactation.

## Acknowledgments

The authors thank the CMI-AII and the Région Bretagne for financial support. The authors are grateful to the staff of the experimental farm of Méjussaume (INRA, Le Rheu, France) and to the staff of the UMR 1348 laboratory (INRA Agrocampus Ouest, Saint Gilles, France) for the laboratory analyses.

## Author Contributions

**Conceptualization**: PG, KL, AL, CH, AB

**Data Curation**: PG, AB

**Formal Analysis**: PG

**Funding Acquisition**: KL, AL, CH, AB

**Investigation**: PG, CH, AB

**Methodology**: PG, CH, AB

**Project Administration**: PG, AB

**Resources**: CH, AB

**Software**: PG

**Supervision**: AB

**Validation**: PG, CH, AB

**Visualization**: PG, CH, AB

**Writing – Original Draft Preparation**: PG, CH, AB

**Writing – Review & Editing**: PG, KL, AL, CH, AB

## Supporting information

**S1 Table. Estimated adjusted means (± Standard errors of the means) of all studied variables according to the treatment and the week of lactation.**

